# Robust Encoding of Abstract Rules by Distinct Neuronal Populations in Primate Visual Cortex

**DOI:** 10.1101/2020.10.28.351460

**Authors:** Donatas Jonikaitis, Nir Nissim, Ruobing Xia, Tirin Moore

**Affiliations:** Department of Neurobiology and Howard Hughes Medical Institute, Stanford University School of Medicine, Stanford, CA 94305, USA; Department of Industrial Engineering and Management, Ben-Gurion University of the Negev, Beer-Sheva 84105, Israel; Malware Lab, Cyber Security Research Center, Ben-Gurion University of the Negev, Beer-Sheva 84105, Israel

## Abstract

It is widely known that neural activity in sensory representations is modulated by cognitive factors such as attention, reward value and working memory. In such cases, sensory responses are found to reflect a selection of the specific sensory information needed to achieve behavioral goals. In contrast, more abstract behavioral constraints that do not involve stimulus selection, such as task rules, are thought to be encoded by neurons at later stages. We show that information about abstract rules is encoded by neurons in primate visual cortex in the absence of sensory stimulation. Furthermore, we show that rule information is greatest among neurons with the least visual activity and the weakest coupling to local neuronal networks. Our results identify rule-specific signals within a sensory representation and suggest that distinct mechanisms exist there for mapping rule information onto sensory guided decisions.

## Results

Many behaviors can rely on fixed stimulus-response associations. However, more complex behaviors, particularly those of primates, rely on the ability to flexibly assign different behavioral responses to particular stimuli, depending on the context. For example, orienting toward a stimulus under one rule, yet avoiding the stimulus under a different rule requires flexible command of behavioral output, and it is a hallmark of executive control (Jonikaitis, Dhawan, & Deubel, 2019; Miller & Cohen, 2001; Munoz & Everling, 2004) The neural mechanisms of behavioral flexibility have been studied extensively in both human and nonhuman primate models, where a dominant role of prefrontal cortex in the implementation of abstract rules is well-established (Fuster, 1997; Mansouri, Freedman, & Buckley, 2020; Passingham, 1993). For example, neurons in prefrontal cortex exhibit robust representations of the abstract rules governing the mapping of sensory stimuli onto behavioral responses (Wallis, Anderson, & Miller, 2001; White & Wise, 1999). A number of studies demonstrate a direct influence of premotor and prefrontal cortex on signals within sensory cortices (Petreanu et al., 2012; Squire, Noudoost, Schafer, & Moore, 2013; Zagha, Casale, Sachdev, McGinley, & McCormick, 2013; Zhang et al., 2014), thus raising the possibility that representations of abstract rules may be propagated to sensory areas. However, the extent to which neurons in sensory areas signal the rules governing stimulus selection, rather than only the selection itself, remains an open question.

It is known that sensory activity within primate visual cortex is influenced by cognitive factors such as attention (Noudoost, Chang, Steinmetz, & Moore, 2010; Reynolds & Chelazzi, 2004), reward expectation (Baruni, Lau, & Salzman, 2015; Stănişor, van der Togt, Pennartz, & Roelfsema, 2013) and working memory (Mendoza-halliday, Torres, & Martinez-trujillo, 2014; Supèr, Spekreijse, & Lamme, 2001a; van Kerkoerle, Self, & Roelfsema, 2017). Typically, variation in these factors is found to influence the selection of specific information by visually responsive neurons, information such as visual stimulus features (Bichot, Rossi, & Desimone, 2005; Motter, 1994) or location (Motter, 1993). This modulation is generally thought to contribute to changes in stimulus related aspects of behavioral performance, such as perceptual sensitivity to attended stimuli (Noudoost et al., 2010; Reynolds & Chelazzi, 2004), reinforcement of particular features (Baruni et al., 2015; Stănişor et al., 2013), or memory of relevant locations (Mendoza-halliday et al., 2014; Supèr et al., 2001a; van Kerkoerle et al., 2017). In such cases, the selection of stimulus information is inherent in a given behavioral condition, and the involvement of sensory neurons in signaling selected stimuli seems clear. Much less clear is whether information about abstract rules is also represented in sensory representations. Often, the rule governing performance on a particular task can be abstract, and thus orthogonal to any particular dimension of sensory input and may merely identify appropriate mappings between sensory input and behavioral responses (Mansouri et al., 2020; Miller & Cohen, 2001). In such cases, neural activity in sensory areas may not reflect abstract rule *per se*. However, this has not been thoroughly examined. One study found that in comparison to premotor and prefrontal cortex, where a high proportion of neurons signaled abstract rules, only very few neurons signaled task rules within inferotemporal cortex, the final stage of the ventral visual system (Muhammad, Wallis, & Miller, 2006). This observation is consistent with the notion that these signals emerge largely outside of sensory representations (Fuster, 1997).

We trained two monkeys (AQ and HB) to perform two versions of a task in which they either looked at, or avoided looking at, a memorized location (Methods and Fig. 1A). In the ‘Look’ task, monkeys memorized a cue location and after a delay period, were rewarded for making an eye movement to a target appearing at the cued location. In the ‘Avoid’ task, monkeys were instead rewarded for making an eye movement to a target appearing at a novel location. Although the behavioral responses differed between the two tasks, looking at or avoiding the cued location, neither could be solved without memorizing that location. The two tasks are similar to the match and non-match memory tasks used to measure abstract rule signals in prefrontal cortex (Mansouri et al., 2020). Monkeys performed alternating blocks of Look and Avoid trials during each experimental session (AQ: 7 sessions; HB: 8 sessions; Figure 1B). Both monkeys performed the two tasks successfully (median performance Look: AQ = 89.6%, HB = 72.7%; median performance Avoid: AQ = 82.5%, HB = 70.2%), and thus could switch between the two task rules. Across sessions, performance varied, and on a given session, could be greater either during the Look or Avoid task. Overall, although average performance was greater in the Look task, this difference was only significant in one monkey (AQ: 6.9%, *p*=0.25; HB: 2.5%, *p*=0.05).

**Fig. 1.**
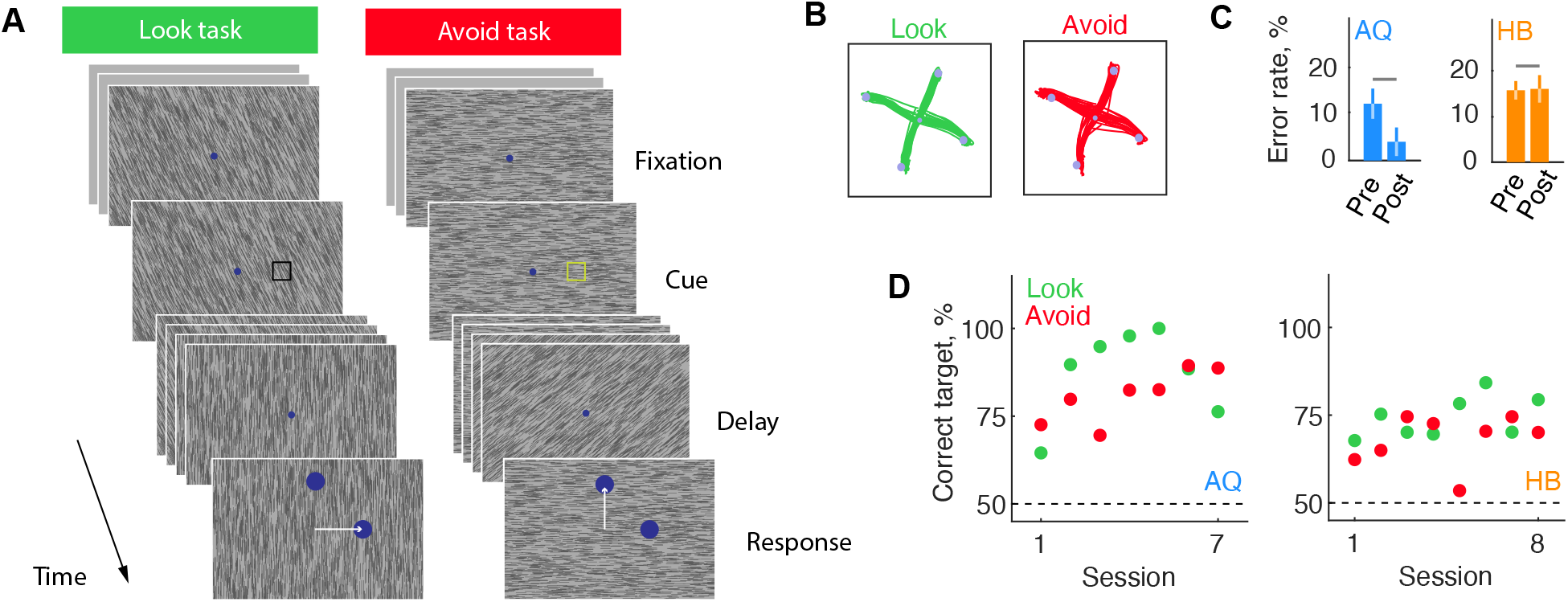
Look and Avoid tasks. **A.** In both tasks, following central fixation, a briefly presented cue (open square) indicated the location to be memorized. After a delay period, two response targets appeared, one at the cued location and one at a location randomly selected from the non-cued locations. In the Look task (left), monkeys made a saccadic eye movement to the target matching the cued location. In the Avoid task, monkeys made an eye movement to the target at the non-matching location. The Look and Avoid tasks were performed in alternating blocks of 150-400 trials. The cue color differed between the different tasks to highlight the current rule. On a majority of trials, the display contained a task-irrelevant background texture; on the remainder of trials, it was uniform gray. **B.** example eye movement responses to four target locations during a single session for Look and Avoid blocks (monkey AQ). **C.** The mean error rates during the last 5 (pre) and first 5 trials (post) relative to task switches (Look to Avoid or Avoid to Look). **D.** Mean performance on each of the two tasks across sessions for the two monkeys.

We asked whether neurons in extrastriate area V4 encoded information about the two tasks prior to the start of a behavioral trial. We focused on a 300-ms, pre-cue period during each task, when only the task rule was known. Neuronal activity was recorded from neurons in area V4 using linear array electrodes which provided simultaneous recordings at 16-24 sites distributed across the cortical depth (Methods and Fig. S1). Single and multi-unit activity were combined totaling 310 units across 15 sessions in the two monkeys (AQ: 7 sessions, 144 units; HB: 8 session, 168 units). Using these data, we first looked for differences in overall firing rate during the pre-cue period. In some sessions, indeed we observed clear differences in pre-cue firing rate among simultaneously recorded neurons between the two tasks. Yet, in other sessions, those differences were absent (Fig. 2A). Across all neurons and sessions, activity was significantly higher in the Look task in both monkeys (Figure 2B; AQ, median modulation index = 0.06 ± 0.014, *p*<0.001; HB, median modulation index = 0.03 ± 0.004, *p*<0.001), amounting to a ~6-12% increase in activity. Notably, the task modulation was evident whether or not neurons were visually driven by a task-irrelevant background texture (Supèr et al., 2001a) (Fig. 2C). However, the change in activity was highly variable across experimental sessions in both monkeys. Therefore, we focused all subsequent analyses on the session-by-session data.

**Fig. 2.**
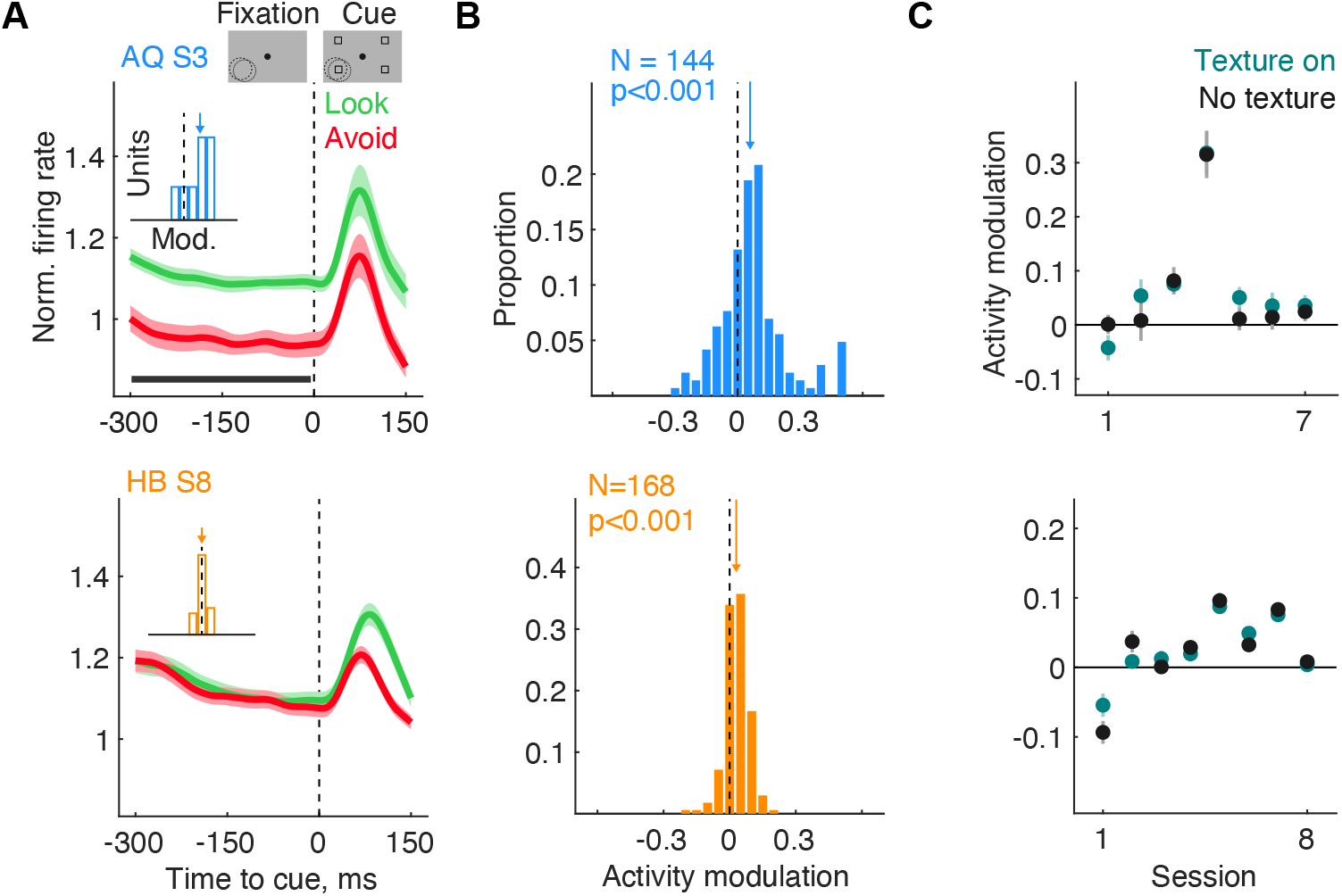
Modulation of V4 neuronal activity during the pre-cue period. **A.** Mean firing rate at 16 simultaneously recorded V4 sites measured during the pre-cue period in an example session from each monkey (top and bottom). Look and Avoid firing rates are averaged across all cue locations. Solid line indicates the period used for pre-cue period data analyses. Inset shows the distribution of activity modulation indices calculated for each unit during an example session. **B.** Distributions of Look-Avoid activity modulation indices for all recorded units in each monkey. **C.** Mean activity modulation indices for each recorded session in each monkey. Data are separated by Texture-on and No-texture conditions.

In other forms of visual cortical modulation, different behavioral conditions tend to produce largely unidirectional changes in activity across all visual cortical neurons. For example, firing rates generally increase for all neurons when attention (Noudoost et al., 2010) or working memory (Supèr et al., 2001a) is directed into a RF, or when RF stimuli have a higher reward expectation (Baruni et al., 2015). However, we considered that rule modulation might behave differently, and that merely comparing mean firing rates could obscure more robust effects of task rule on the pattern of spiking activity. Thus, we adopted a decoding approach to further examine how robustly task rule was encoded by the activity of V4 neurons. Using the same data set, we employed six commonly used machine learning algorithms to measure the extent to which task rule could be accurately decoded during the pre-cue period from the activity of neurons simultaneously recorded during each session (Fig. 3A). The chosen algorithms comprise a broad set of complementary approaches based on different decoding principles (Methods). In each experimental session, we measured the accuracy of each decoder in classifying the task rule (Look/Avoid) using the spiking activity of simultaneously recorded neurons. Remarkably, we found that in both monkeys, each of the six decoders could accurately classify the task above chance level (Fig. 3B; AQ: 83-94%, HB: 68-79%). Moreover, we observed this in every experimental session (Fig. 3C; AQ for each session p<0.001; HB for each session p<0.001). Notably, decoding accuracy was unrelated to any differences in behavioral performance observed between the two tasks (Fig. S2). Furthermore, decoding accuracy was statistically equal with or without the background texture stimulus, indicating that visually driven activity did not contribute to the encoding of task rules (Fig. 3D; AQ: ANOVA F(1,6) = 3.79, *p*=0.1; HB: F(1,7) = 4.57, *p*=0.07). Thus, in spite of the variability in overall firing rate effects across sessions, rule information was invariably present in V4 neuronal activity.

**Fig. 3.**
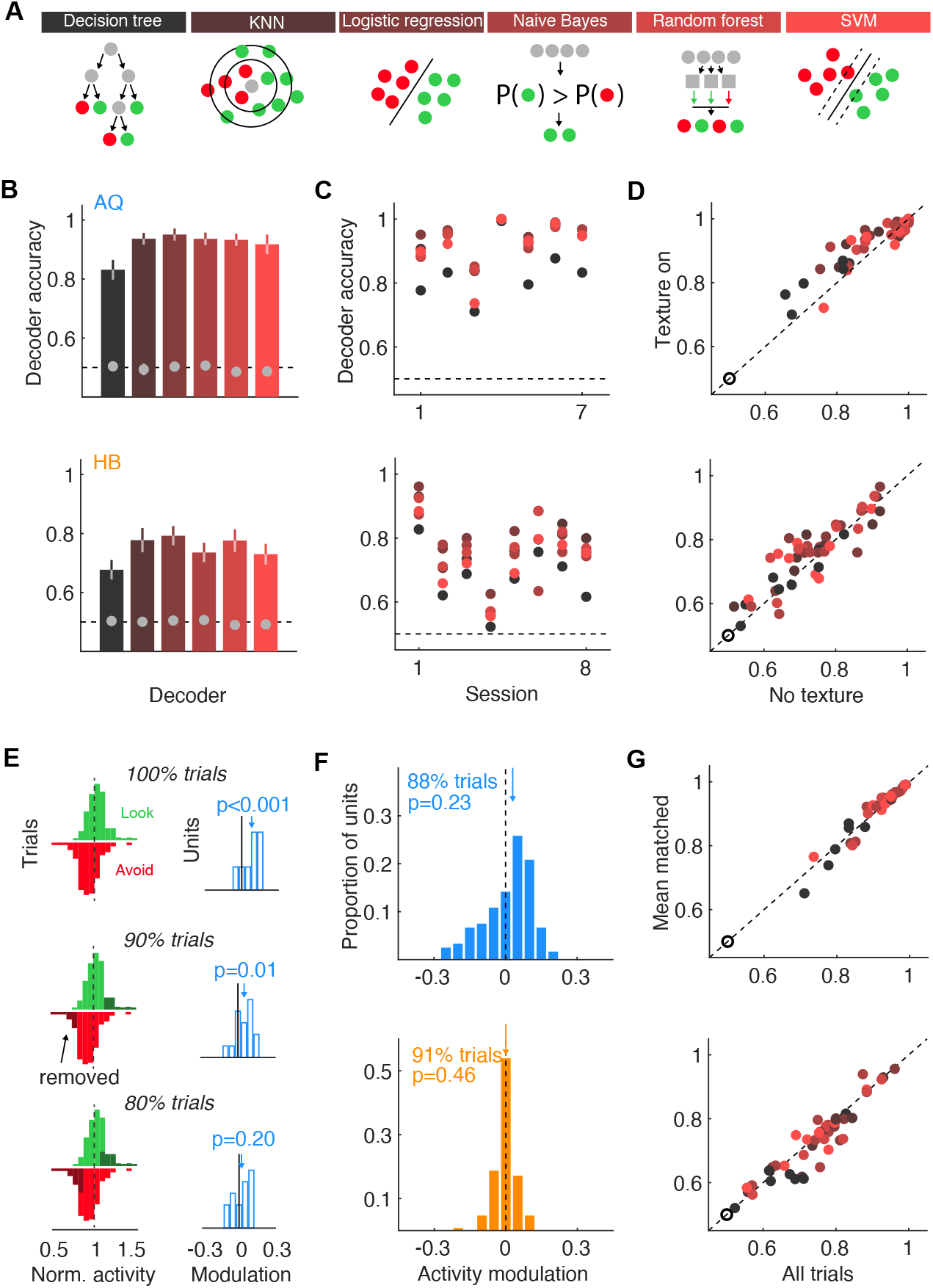
Decoding of task rule from V4 neuronal activity during the pre-cue period. **A.** Six different complementary machine learning algorithms were used to decode task rule from V4 activity. Diagrams shown with each algorithm highlight the differences between decoding approaches. **B.** Mean decoding accuracy for each algorithm across all sessions in both monkeys (top, bottom). Filled circles denote accuracy obtained from label-shuffling. **C.** Decoding accuracy for each individual session and each decoding algorithm for both monkeys. Dashed line indicates chance performance (accuracy = 0.5). **D.** Decoding accuracy for Texture-on versus No-texture trials in both monkeys. **E.** Mean-matching procedure for an example session (AQ session 3; same data as Figure 2A). Activity modulation when all data (100^th^ percentile) are used (top). Left panel shows distributions of normalized activity (green and red histograms) calculated across all units across trials. In this session, mean activity in the Look task exceeded that of the Avoid task. Right histogram shows the distribution of activity modulation indices across recorded units in the session. In this example session, modulation was still significant after removal of the upper and lower 10% of Look and Avoid trials, respectively (90^th^ percentile). Modulation was nonsignificant for the 80^th^ percentile trials. **F.** Distributions of Look-Avoid activity modulation indices for all recorded units in each monkey following mean-matching. **G.** Decoding accuracy for all trials compared to mean-matched trials. All error bars denote standard errors of the mean. Some error bars are smaller than the symbols.

The fact that differences in neuronal activity between the two tasks could be detected by the decoders in every experimental session suggests that decoder performance was driven by factors independent of overall mean firing rate. We confirmed this directly by measuring decoding accuracy in subsets of trials in which the mean firing between the two tasks was reduced or eliminated. In datasets obtained from each session, we excluded trials on which overall mean activity in Look and Avoid trials deviated most from the overall mean (Fig. 3E). As a result of this mean-matching procedure, 88% - 91% of the total trials remained in the analysis, and the average task modulation was reduced below statistical significance in both monkeys (Fig. 3F; AQ: median modulation = 0.03±0.01, *p*=0.23; HB: median modulation = 0±0.001, *p*=0.46). In spite of the removal of overall mean firing rate differences between the two tasks, decoding of task rule remained accurate in every session of data from both monkeys (Fig. 3G). Moreover, when comparing the accuracies of decoding performance, we observed no significant differences in the mean-matched trials and all trials (AQ: ANOVA, F(1,5) = 0.00, *p*=0.95; HB: F(1,7) = 1.38, *p*=0.28). This result demonstrates that rule information is largely encoded in the pattern of spiking activity across neuronal populations in V4, rather than in unidirectional firing rate changes, as classically observed in visual cortex.

In addition to cognitive effects, previous studies have also shown that changes in arousal state can influence visual cortical activity (Beaman, Eagleman, & Dragoi, 2017; Engel et al., 2016). Thus, we considered that switches in task rule might systematically affect arousal (e.g. increase during the Avoid task), and in turn might alter visual cortical activity. Pupil size has been widely used as an index of arousal (Mathôt & Van der Stigchel, 2015; Reimer et al., 2014). So, we compared pupil size between the two tasks (Fig. S3A). However, we found no consistent differences in pupil size across the two tasks in either monkey, suggesting that arousal was largely equal during performance of the two tasks. Moreover, we observed no systematic effect of previous reward outcome on pupil size in either task (Fig. S3B). In addition, as it is known that fixational eye movements (microsaccades) can alter visual cortical activity (Leopold & Logothetis, 1998; Lowet et al., 2018), we also compared microsaccadic behavior between the two tasks. Again, we found no consistent differences in the pattern of microsaccades between the two tasks (Fig. S4). Moreover, since the effects of task rule on activity were observed even in the absence of visual stimulation, differences in microsaccadic behavior would not be expected to influence visually driven activity. Lastly, we considered whether pre-cue activity might reflect a representation of the anticipated cue stimulus. However, consistent with previous studies demonstrating a lack of feature-based effects in V4 in the absence of visual stimulation (Chelazzi, Miller, Duncan, & Desimone, 2001), we found no evidence that neurons signaled the upcoming cue stimulus (Fig. S5).

A number of previous studies have found evidence that subpopulations of neurons within visual cortex may be engaged differently during attention (Mitchell, Sundberg, & Reynolds, 2007; Nandy, Nassi, & Reynolds, 2017; Pettine, Steinmetz, & Moore, 2019) and working memory (van Kerkoerle et al., 2017). Thus, we next asked if the rule signals we observed might be driven disproportionately by distinct subsets of neurons recorded in the daily sessions. To test that possibility, we took advantage of the ‘feature importance’ coefficients yielded from the decoding analyses; these coefficients scale the influence of each of the component neurons in the population on the decoder’s performance (Golub et al., 1999; Guyon & Elisseeff, 2003) (Methods). As expected, in each session, the distributions of feature importance were broad (Fig. S6), indicating that particular neurons contributed disproportionately to the encoding of task rule. Importantly, feature importance was strongly correlated with the absolute value of the task modulation indices computed across neurons in both monkeys (Fig. S7). Using this information, we first sought to determine the relationship between rule information and the visual responsiveness of V4 neurons. Since attentional modulation tends to co-occur with the largest visual responses (Bichot et al., 2005; McAdams & Maunsell, 1999; Motter, 1993; Noudoost et al., 2010; Reynolds, Pasternak, & Desimone, 2000), we considered that feature importance might be largest among neurons exhibiting the largest visual responses. However, to our surprise, we found the opposite in the session-by-session data in both monkeys (Fig 4A-B). Task rule information (feature importance) was negatively correlated with the magnitude of visual response magnitude across neurons (Fig. 4C) (AQ, mean r = −0.3, *p*=0.03; HB, mean r = −0.35, *p*<0.001; combined mean r = −0.33, *p*<0.001), indicating that neurons conveying the most information about task rule were those with the least visually evoked activity. A recent study revealed that in both mouse and primate visual cortex, neurons vary broadly in the extent to which their activity is coupled to the population firing rate (Okun et al., 2015). Importantly, the data show that population coupling predicts the magnitude of visually evoked activity and of attention-related modulation; strongly coupled neurons exhibit the largest visual responses and the largest modulation. Given our observation of reduced visual activity among neurons signaling task rule, we considered that the most informative neurons might correspond to subpopulations with weak population coupling. Thus, we measured population coupling for all of the recorded neurons using the spike-triggered average population response for each unit and tested this hypothesis (Methods and Fig. 4D). As in the previous study, population coupling was positively correlated with the magnitude of visually driven activity, a pattern that was observed across experimental sessions and in both monkeys (Fig. 4E-G; AQ, mean r = 0.64, *p*<0.001; HB, mean r = 0.39, *p*=0.007; combined mean r = 0.51, *p*<0.001). However, we also found that population coupling was negatively correlated with feature importance, a pattern that was also observed across sessions and monkeys (Fig. 4H-J; AQ, mean r = −0.32, *p*<0.001; HB, mean r = −0.28, *p*=0.04; combined mean r = −0.3, *p*<0.001). Thus, neurons conveying the most information about task rules comprised a distinct subpopulation of cells with the least visual activity and weak coupling to the local population.

**Fig. 4.**
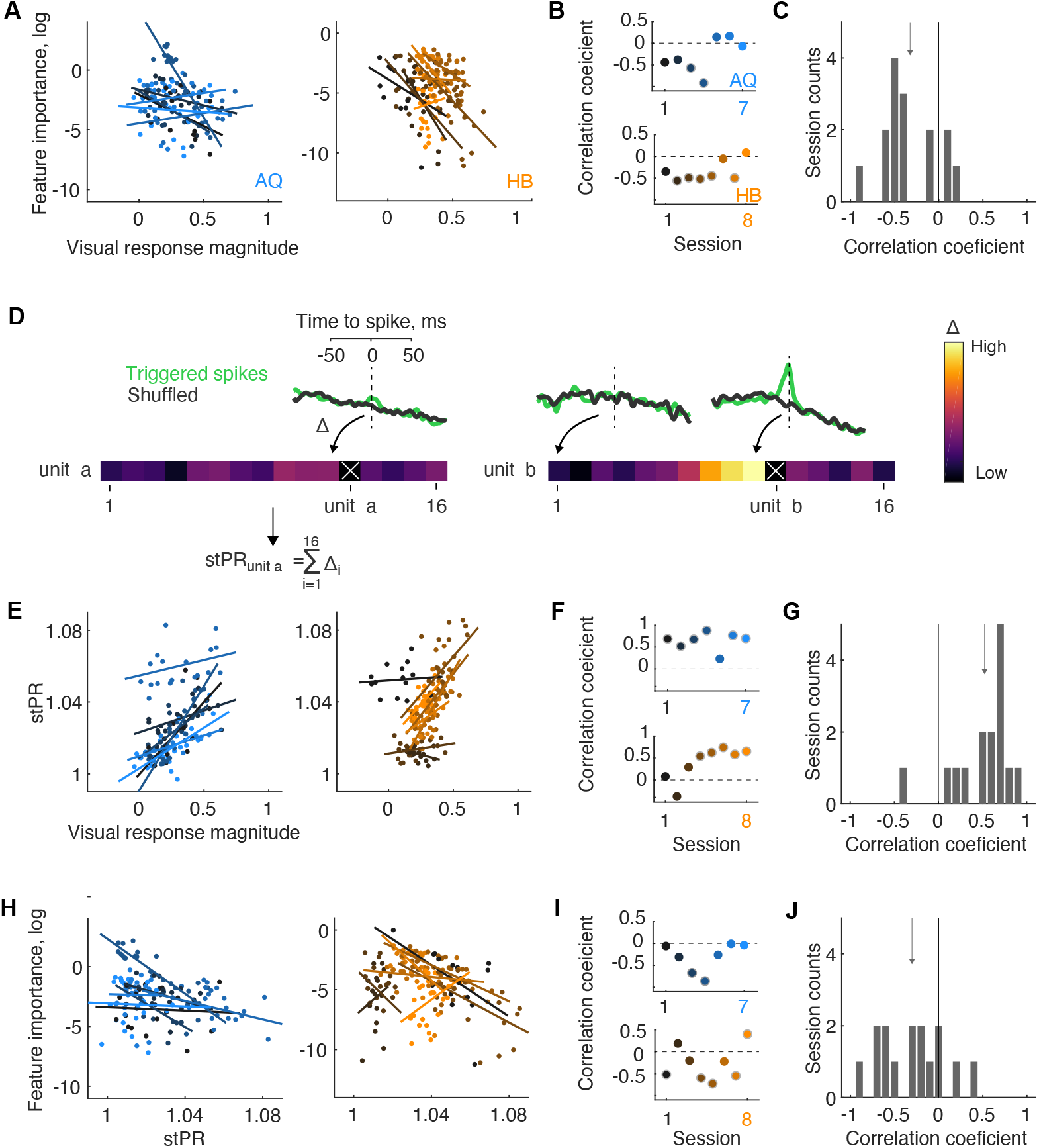
Relationship of feature importance to visual activity and population coupling. **A.** Correlations between visual response magnitude and feature importance for each session and each monkey. Each set of points and its linear fit shows the trade-off between neuronal feature importance in the decoder and the magnitude of each neuron’s response to the visual cue. **B.** Correlation coefficients obtained for each session and each monkey. **C.** Distribution of correlation coefficients combined across both monkeys. **D.** Computing the stPR in two example units (a) (left) and b (right). Population rates were first computed from the accumulation of spikes from all units except the unit in question (indicated by ‘X’). The stPR was then computed as the spike rate triggered on the spikes from that unit (green line). In the two examples, the stPR is greater in b than a. Black line shows the spiketime shuffled stPR. **E-G.** Correlations between visual response magnitude and the stPR across sessions in each monkey. **H-J.** Correlations between feature importance and stPR.

Lastly, we found that neurons with robust task rule signals were distributed equally across layers of V4. Previous studies have shown that both attention and working memory influence neuronal activity differently across layers of visual cortex, including in V4 (Nandy et al., 2017; Pettine et al., 2019; van Kerkoerle et al., 2017). Since our array recordings were made largely perpendicular to the cortical surface (Fig. S1, S8A), we could assess whether rule related signals were distributed differently across cortical layers. As in our previous analyses, we looked for laminar differences in the session-by-session data. These comparisons revealed that the contribution of neurons to rule information was distributed equally across cortical layers (Fig. S8B). In addition, rule information did not vary significantly with distance from input layer 4 (AQ: r = −0.01, *p*=0.89; HB: r = 0.11, *p*=0.19) (Fig. S8C). Thus, although task rule was encoded by a distinct subpopulation of neurons, those neurons were evenly distributed across cortical layers.

How generally might visual cortical neurons encode abstract rules? Although the two rules we employed were sufficient to produce robustly decodable signals in V4, it need not follow that V4 neurons signal differences between any arbitrary set of rules. In contrast to prefrontal cortex, where neurons appear to signal a multitude of arbitrary stimulus-response mappings (Mansouri et al., 2020; Miller & Cohen, 2001), one would not necessarily expect the same of visual cortical neurons. Rather, one might expect rule-related signals to emerge only in tasks that draw upon the unique functions of visual cortical circuitry, including perhaps non-sensory functions. Consistent with that notion, the modulation of visual cortical activity observed by attention, reward or working memory are typically interpreted as reflecting the role of visual cortical neurons in signaling the perceptual relevance of associated sensory stimuli (Baruni et al., 2015; Reynolds & Chelazzi, 2004; Supèr et al., 2001a). In contrast, the modulation we observed indicates that visual cortical activity also signals how forthcoming input is to be mapped onto subsequent behavioral responses. On the one hand, this observation is at odds with the classical view that visual cortical areas function largely as passive, perceptual filters, particularly areas within the primate ventral stream (Milner & Goodale, 2006), such as area V4. On the other hand, evidence of a more active role of visual cortical neurons in eye movement preparation has been accumulating for some time (Moore, Armstrong, & Fallah, 2003; Steinmetz & Moore, 2014). Thus, the observation that V4 neurons also signal rules about how visual stimuli are to be mapped onto eye movements might reflect an engagement of visual cortical activity in the preparation of appropriate oculomotor responses. In primates, prefrontal connections with visual cortex occurs largely via direct connections between gaze control neurons within the frontal eye field and extrastriate areas (Merrikhi et al., 2017; Stanton, Bruce, & Goldberg, 1995). These connections might provide one source of the activity differences we observed between tasks involving different oculomotor strategies, though it remains to be determined.

A number of studies have focused on identifying the circuits underlying changes in visual cortical processing resulting from extra-retinal factors, including in primates (Fiebelkorn & Kastner, 2020; Krauzlis, Lovejoy, & Zénon, 2013; Noudoost et al., 2010). Evidence from these studies reveals a number of specific circuits each capable of altering visual cortical activity, including input from thalamus (Saalmann, Pinsk, Wang, Li, & Kastner, 2012), prefrontal cortex (Squire et al., 2013), or from diffuse neuromodulatory nuclei (Herrero et al., 2008). Although these mechanisms are often invoked as possible sources of attention-related modulation, their respective roles in other functions, such as reward and working memory remain largely unexplored. It is possible that each modulatory mechanism contributes to a specific function, but it is also possible that each mechanism contributes some complementary aspect to multiple functions. The rule signals we describe here thus far appear distinct from previously described modulations of visual cortical activity. Their prevalence among neurons with minimal visual responsiveness differs from the effects of attention (Bichot et al., 2005; McAdams & Maunsell, 1999; Motter, 1993; Noudoost et al., 2010; Reynolds et al., 2000), and their prevalence among neurons weakly coupled to the population contrasts with other top-down influences on V4 activity (Okun et al., 2015). Nevertheless, it remains to be seen whether these rule related signals emerge from mechanisms that are themselves distinct from those driving other types of visual cortical modulation.

## Materials and Methods

### General and Surgical Procedures

Two male rhesus monkeys (Macaca mulatta, 10 and 13 kg), monkey AQ and monkey HB, were used in this study. All experimental procedures were in accordance with National Institutes of Health Guide for the Care and Use of Laboratory Animals, the Society for Neuroscience Guidelines and Policies, and Stanford University Animal Care and Use Committee and are detailed in a previous report (Armstrong, Chang, & Moore, 2009).

### Behavior: Look and Avoid tasks

Experiments were controlled by a DELL Precision Tower 3620 desktop computer and implemented in Matlab (MathWorks, Natick, MA, USA) using Psychophysics and Eyelink toolboxes (Brainard, 1997; Cornelissen, Peters, & Palmer, 2002). Eye position was recorded with an SR Research EyeLink 1000 desktop mounted eye-tracker (sampling rate of 1000 Hz). Stimuli were presented at a viewing distance of 60 cm, on an VIEWPixx3D display (1920 x 1080 pixels, vertical refresh rate of 60 Hz). Each behavioral trial began with a fixation spot (a circle of radius 0.5°, degrees of visual angle, dva) presented at the center of the screen (gray background). After the monkey acquired and maintained fixation for 600-800 ms, a cue appeared (colored square frame, size 1° x 1° dva) for ~50 ms at a randomly selected location (1 of 4 possible locations separated by 90° polar angle; eccentricity from fixation ranged from 5° to 7° dva). Cue presentation was followed by a delay period (1600-1800 ms on majority of sessions; as short as 1000-1200 ms and as long as 2000-2200 ms), after which fixation disappeared and the response targets appeared. Monkeys received a juice reward for making a saccadic eye movement to the correct target and maintaining gaze on the target for 200 ms. Failures to acquire fixation, breaks of fixation during the delay, or incorrect eye movements were not rewarded.

#### Look task

The cue (open square) color was black for monkey AQ and green for monkey HB. On 93% of trials, after the delay period, two targets appeared (filled blue circles of radius 1° dva). One of the targets always appeared at the previously cued location while the other appeared at a randomly selected one of the other of the 3 remaining locations. To be rewarded, monkeys had to make a saccadic eye movement to the target at the cued location. On 7% of trials, (probe trials), after the delay, only one target appeared (filled black circle, at cued or 180° opposite), and monkeys had to make a saccadic eye movement to the probe target to be rewarded.

#### Avoid task

Avoid trials were identical to the Look task, except that to be rewarded monkeys had to make a saccadic eye movement to the novel target, i.e. the one not previously cued. The cue color was green for monkey AQ and black for monkey HB. As in the Look task, probe trials occurred on 7% of trials. Look and Avoid task blocks were interleaved. Block duration varied, typically ranging from 150 to 400 trials per block. Each session could begin either with a Look or an Avoid block.

### V4 recording and visual stimuli

Recording sites within area V4 were identified by neuronal responses to visual stimuli and by assessing receptive field sizes as a function of stimulus eccentricity and receptive field location (Gattass, Sousa, & Gross, 1988). Recordings were obtained with 16 or 24-channel linear array electrodes with contacts spaced 75 or 150 m apart (U-Probes and V-Probes, Plexon, Inc).

Electrodes were lowered into the cortex using a hydraulic microdrive (Narishige International) at angles roughly perpendicular to the cortical surface. The receptive fields of V4 neurons studied were located within the lower contralateral quadrant within ~4° to 8° eccentricity. Neuronal activity was measured against a local reference, a stainless guide tube, which was close to the electrode contacts. Data were amplified and recorded using the Omniplex system (Plexon Inc., Dallas, TX). Wide-band data filtered only in hardware at 0.5Hz highpass and 8kHz lowpass, were recorded to disk at 40kHz. Data in each channel was then extracted using 3 standard deviations as a threshold for extracellular spike and classified as multi-unit activity.

During the Look and Avoid tasks, at least one of the four cues appeared within the receptive fields of recorded neurons. In order to evoke activity in the recorded neurons during the pre-cue period, the display background was filled with a task irrelevant texture (Figure 1A) beginning 600 ms before cue onset (Supèr, Spekreijse, & Lamme, 2001b). The background texture consisted of a dense field of 10000 oriented lines (width: 2 pixels, length: 2° dva), one orientation per background, selected randomly from 0° to 179° in 30° increments. The background texture was presented on ~83% of trials; on ~17% of trials, the background remained uniform gray.

### Data analysis

#### Behavior

Gaze position on each trial was drift corrected by using median gaze position from 10 previous trials. Drift correction was based on gaze position from 100 ms to 10 ms before the cue onset, when stable fixation was maintained. We detected saccades offline using an algorithm based on eye velocity changes (Engbert & Kliegl, 2003). We next clustered saccades as ending on one of the three potential locations: (1) fixation, (2) correct response target, (3) wrong response target. The clustering procedure used support vector machine algorithm with a Gaussian kernel (Jonikaitis et al., 2019). Saccades directed to the target or distractor had a latency of at least of 50 ms after the response cue (Fischer & Boch, 1983). For the microsaccade analysis, we used all saccades that did not break the fixation window during the pre-cue period, i.e. saccades with amplitudes less than 1 dva. We removed trials if blinks occurred from 100 ms before cue onset to the time of saccade target onset. Data from each recording was inspected for saccade detection accuracy and data recording noise.

#### Statistical tests

For statistical comparisons of paired-means, we drew (with replacement) 10000 bootstrap samples from the original pair of compared values. We then calculated the difference of these bootstrapped samples and derived two-tailed *p* values from the distribution of these differences. For repeated measures analysis with multiple levels of comparisons (e.g. accuracy of 6 decoders), we used one-way and two-way repeated measures ANOVAS. All correlations were computed as Pearson coefficients. All post-hoc comparisons were based on bootstrap tests and were Bonferonni corrected.

Pre-cue activity was measured during the last 300 ms period prior to the cue presentation. Activity modulation between conditions was calculated using a standard normalized difference: (Look-Avoid)/(Look+Avoid).

Visual responses following the cue onset was measured during a 50-ms window beginning 100 ms after cue onset. This period was chosen as it included the peak of activity in which the response difference between Look and Avoid cue was greatest. Visual response magnitude was defined as (Cue_in – Cue_out) / (Cue_in + Cue_out).

Decoding of mean-matched data: Mean matching was performed for each session. First, for each selected percentile level (95^th^, 90^th^, 85^th^, 80^th^ and 75^th^ percentile), we removed trials with the highest and lowest activity for a given percentile. From this subset of data, we calculated a modulation index across channels (that is, 16 or 24 modulation indices per recording). We then measured whether distribution of those modulation indices differed from 0. If *p*>0.1, this reduced dataset was used as a mean matched data set. For one session we could not mean match the data (AQ session 3). Percentiles for mean matched individual session data were (in order of the sessions): AQ – [90, 90, 80, non-matched, 90, 90, 90]; HB – [90, 100, 95, 95, 80, 90, 75, 100].

### Decoding of V4 activity

We analyzed neuronal spike rates measured during the pre-cue time windows (300 ms). We used six different, commonly used, machine learning algorithms to measure how well pre-cue activity could be used to decode the current task: Look vs. Avoid. Specifically, we used a Decision Tree, Naïve Bayes, K-Nearest Neighbors (KNN), Logistic Regression, Random Forest, and a support vector machines (SVM) algorithm to train classifiers from each experimental session. We chose six standard algorithms, as opposed to one, as a means of corroborating the decoding results using complementary decoding schemes. The Decision Tree algorithm we used is based on the entropy and information gain theorem, by which the algorithm creates its decision paths by prioritizing the features (neurons) whose values (firing rate) provide a maximal amount of new information (AKA Information Gain) when forming the decision tree (Quinlan, 1987). The Naïve Bayes algorithm is a maximum likelihood estimator which is mainly based on Bayes theorem, and conditional and prior probabilities that exist in the data (Duda, Hart, & Stork, 2001). The KNN algorithm does not induce a model (in contrast to the other five algorithms used); instead it is based on distance calculations (e.g., the Euclidian Distance) between each pair of samples (e.g., trials) in the data. A new trial is classified based on the class majority of the K samples to which the trial has maximal resemblance (Cover & Hart, 1967). The logistic regression algorithm is suitable for binary classification problems, like those in our data. To produce an accurate model, it uses a nonlinear (sigmoidal) function probability calculation to select the maximum likelihood fitting curve for the given data, based on an iterative process of data point shifting (Kleinbaum & Klein, 2002). The SVM algorithm is based on solving a quadratic programming optimization problem in an attempt to find a separating hyperplane between the two classes in the given data, so that the distance (i.e., margin) between the samples of the two classes is maximal. We also used the radial basis function kernel (RBF) which projects the given data into a higher dimensional space, thus achieving a better separation of the data and improved decoding accuracy (Cortes & Vapnik, 1995). The random forest algorithm is based on the randomization and selection of different feature subsets and the creation of a sub-decision tree for each subset of features (Ho, 1995). This yields a set (forest) of sub-decision trees, and ultimately an ensemble of decisions and a final decision is produced by using a majority over intermediate decisions.

Each of the six algorithms was trained on all of the datasets, and each classifier employed was evaluated separately on the datasets from each recording session using a standard 10-fold cross-validation procedure where 90% of the dataset was used for training, and the remaining 10% was used for testing. The evaluation included 10 repetitions in which the divisions of the training and test set were randomized with respect to the trials in order to reduce the variance. The session’s classification performance was computed from an area under the curve (AUC) measure and was based on the average AUC of the 10 repetitions. Since the best performance was achieved when using bin size 300 ms (one bin), we only present the performance of decoders for a single bin window.

Feature importance coefficients were obtained by applying a *feature selection* method using a *filters* approach (Guyon & Elisseeff, 2003) in which a specific measure is used to evaluate the correlation of each individual feature (i.e. neuron) to the class (Look/Avoid). A variety of measures and methods exist, but we chose to use the simple, yet efficient, *Fisher score* method (Golub et al., 1999). This method measures the difference between the negative (i.e. Avoid) and positive (i.e Look) samples (i.e. trials) relative to a certain feature (i.e. neuron), in terms of mean and standard deviation. Equation 1 defines the calculation applied by the Fisher score method for each feature, in which *Ri* is the calculated rank of a specific feature *i* (neuron *i*), demonstrating the proportion of the substitution of the mean of the feature *i* values in the negative examples (*N*) and the positive examples (*P*), and the sum of the standard deviation. The larger the calculated *Ri* value, the larger the difference between the values of positive and negative examples relative to feature *i*. Thus, a higher value feature is more important for discriminating between the Avoid and Look trials.

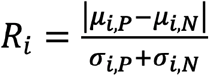

After applying the Fischer score feature selection method on neuronal activity, each neuron was also given a rank. The rank quantified each neuron’s expected contribution to the classification task (regardless of the machine learning algorithm) from which neurons with the top *X* ranks can be selected to train the learning algorithm and yield an accurate classifier.

### Spike-triggered population response. Population coupling

For each unit, a spike-triggered population response (stPR) was computed similar to methods described previously (Okun et al., 2015). The stPR captures the relationship of a given unit’s activity to the summed activity of all neurons recorded in the recorded population at a given moment in time (Fig. 4D). In computing the stPR, the population rate was first measured by accumulating all the detected spikes in the interval −100 to 100 relative to spike onset with 1 ms resolution, and smoothing the result with a Gaussian of half-width 12 ms. For each unit in question, this rate did not include the spikes of that unit and was used to compute the unit’s stPR. Each unit’s stPR was normalized by the time-shuffled stPR.

### Laminar designation

To identify the input layer, we performed a current source density (CSD) analysis for each recording session (Higley, 2012). LFP signals were aligned to the onset of visual cue and averaged over all trials to produce the event related potential (ERP) of each recording channel. The CSD is commonly estimated as the second order derivative of ERPs across channels (Mitzdorf, 1985). To reduce the effect of channels with poor signal quality and obtain a smoother CSD pattern, we conducted an optimization procedure based on the assumption that CSD is a smooth function across cortical depth, which led to the following constrains: 1) the third derivative of ERPs is small, and 2) the modified ERPs do not deviate much from the original data. After the optimization procedure, we calculated the second order derivative of the modified ERPs. For each session, the center depth of the input layer was then identified as the channel that showed the earliest sink in the CSD pattern. Based on the input layer designation, we could compare different layers across cortical depths.

## Acknowledgments

We thank Xiaomo Chen and Joni Wallis for helpful comments on earlier versions of this manuscript and Danielle A. Lopes for technical assistance.

## Funding

This work was supported by NIH Grant EY014924.

## Competing interests

Authors declare no competing interests.

## Data and materials availability

We report sample size, all data exclusions, all data manipulations, and all measures in this study. Summarized experimental data, analysis code and extended set of figures will be available online after publication on zenodo.

---

**Fig. S1.**
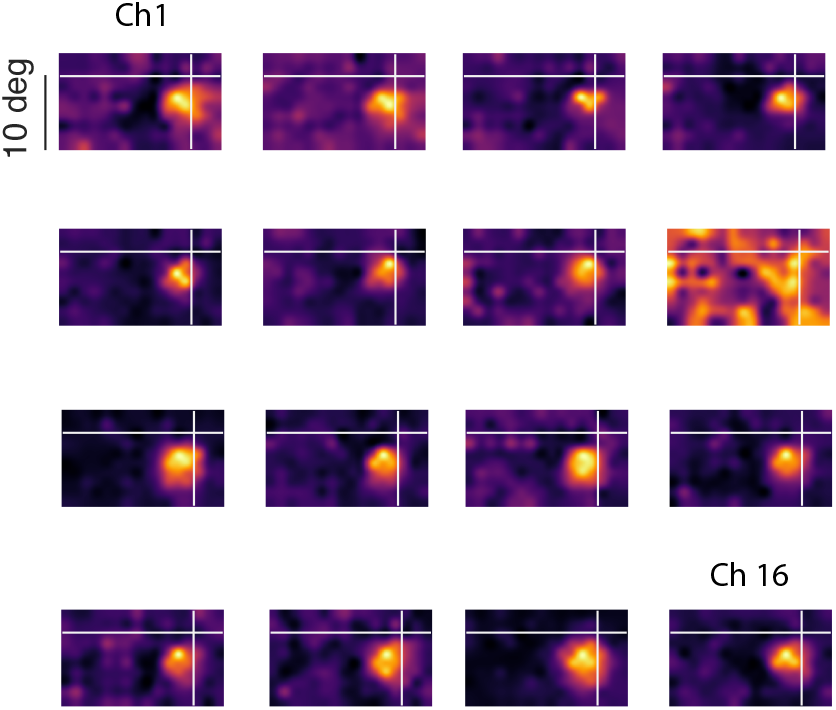
An example linear array recording showing neuronal RFs measured across 16 recording sites spaced every 150 μm across the cortical depth (2250 μm total). In this case, successive, largely overlapping RFs were obtained in all, but one of the 16 recording sites.

**Fig. S2.**
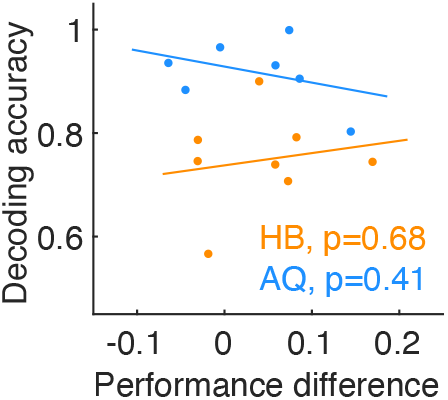
Relationship of task performance (difference between Look and Avoid tasks) to decoding accuracy.

**Fig. S3.**
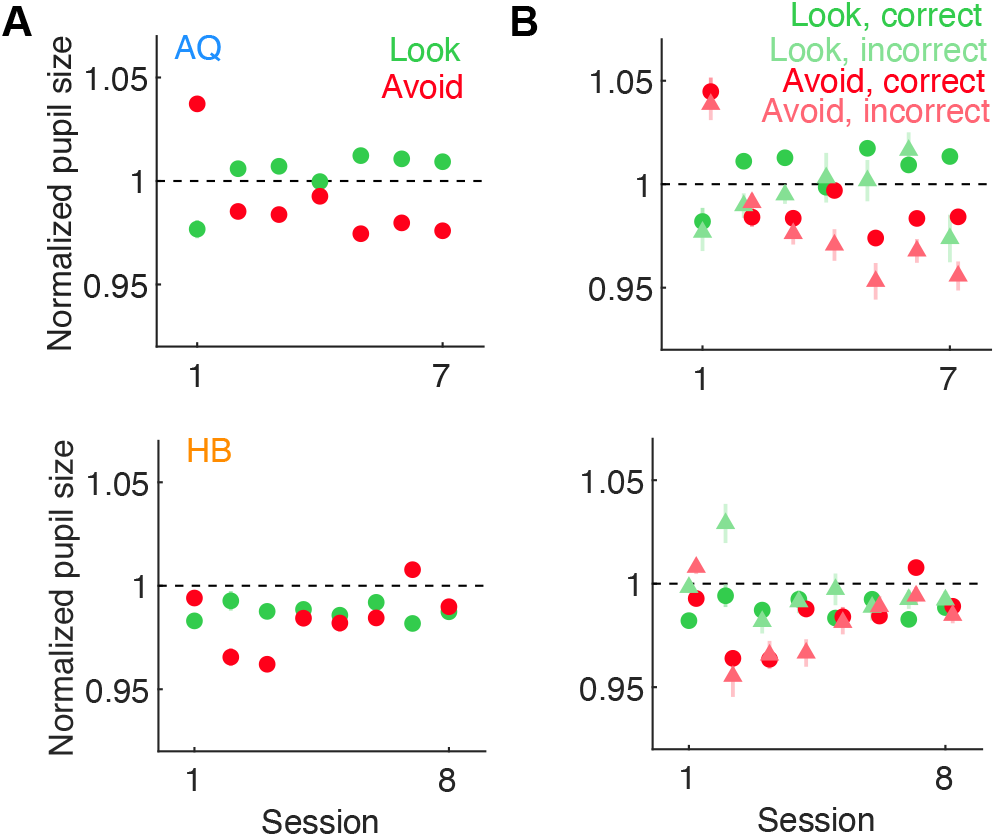
Comparison of pupil size across Look and Avoid tasks in both monkeys. **A.** Pupil size in each session is shown normalized to the session mean. Small differences in pupil size between the Look and Avoid tasks were observable in some sessions. However, those differences were not consistent across experimental sessions in either monkey (AQ: Look = 1 ± 0.005, Avoid = 0.99 ± 0.008, *p*=0.28; HB: Look = 0.99 ± 0.001, Avoid = 0.98 ± 0.005, *p*=0.53). **B.** The influence of previous trial outcome (i.e. reward, no reward) on pupil size across Look and Avoid tasks in the 2 monkeys. A small reduction in pupil size was observed after incorrect trials in one monkey (AQ, F(1,6) = 11.56, *p*=0.01, HB: F(1,7) = 0.62, *p*=0.46). However, there was no effect of task on pupil size (AQ: F(1,6) = 1.09, *p*=0.33, HB: F(1,7) = 1.83, *p*=0.21) and no significant interaction between task and trial outcome in either monkey (AQ: F(1,6) = 0.03, *p*=0.86, HB: F(1,7) = 3.80, *p*=0.09).

**Fig. S4.**
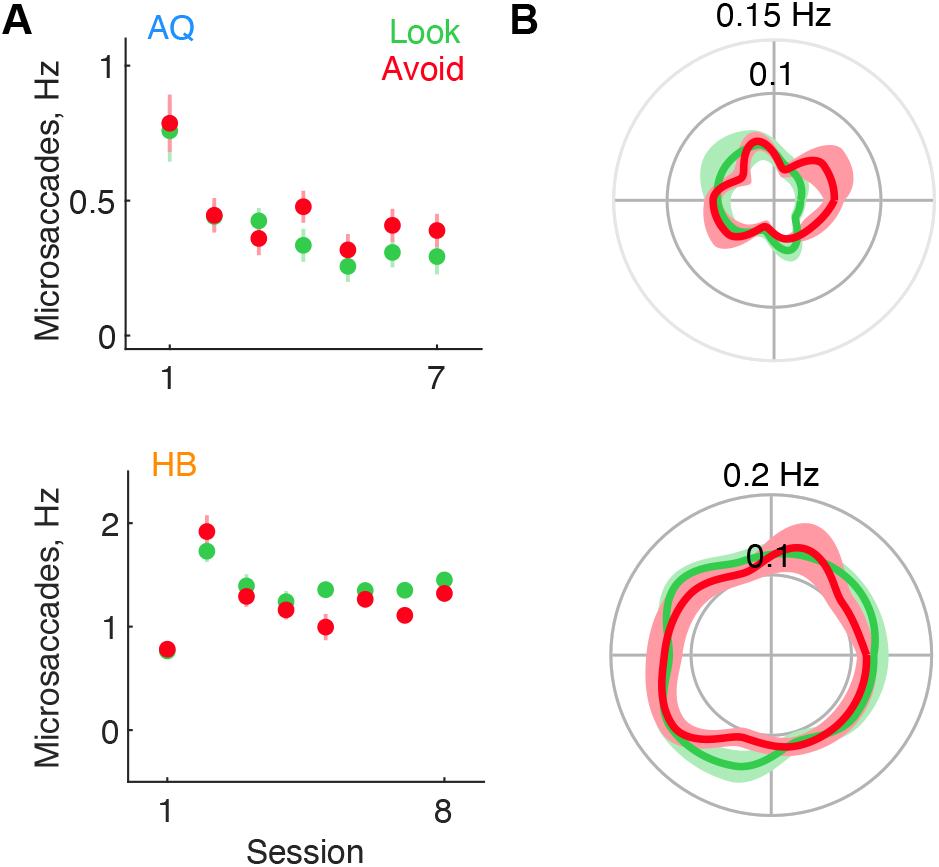
Comparison of fixational eye movements (microsaccades) between the Look and Avoid tasks. **A.** The frequency of microsaccades in the 2 tasks across sessions in the 2 monkeys. The overall frequency of microsaccades in monkey AQ was slightly, but significantly, greater during Avoid trials (mean Look = 0.4 ± 0.07, mean Avoid = 0.45 ± 0.06 Hz, *p*=0.04). However, this pattern was reversed in monkey HB (mean Look = 1.33 ± 0.1, mean Avoid = 1.23 ± 0.11 Hz, *p*=0.06). **B.** The distribution of microsaccade frequencies across movement directions. In both monkeys, the distributions of microsaccade frequencies were statistically indistinguishable across directions (AQ, F(9,54) = 1.52, *p*=0.16; HB, F(9,63) = 1.14, *p*=0.34) and the two tasks (AQ: F(1,6) = 5.32, *p*=0.06; HB: task F(1,7) = 2.85, *p*=0.14) and there was no significant interaction (AQ: F(9,54) = 1.38, p=0.22; HB: F(9,63) = 340.87, *p*=0.56).

**Figure S5.**
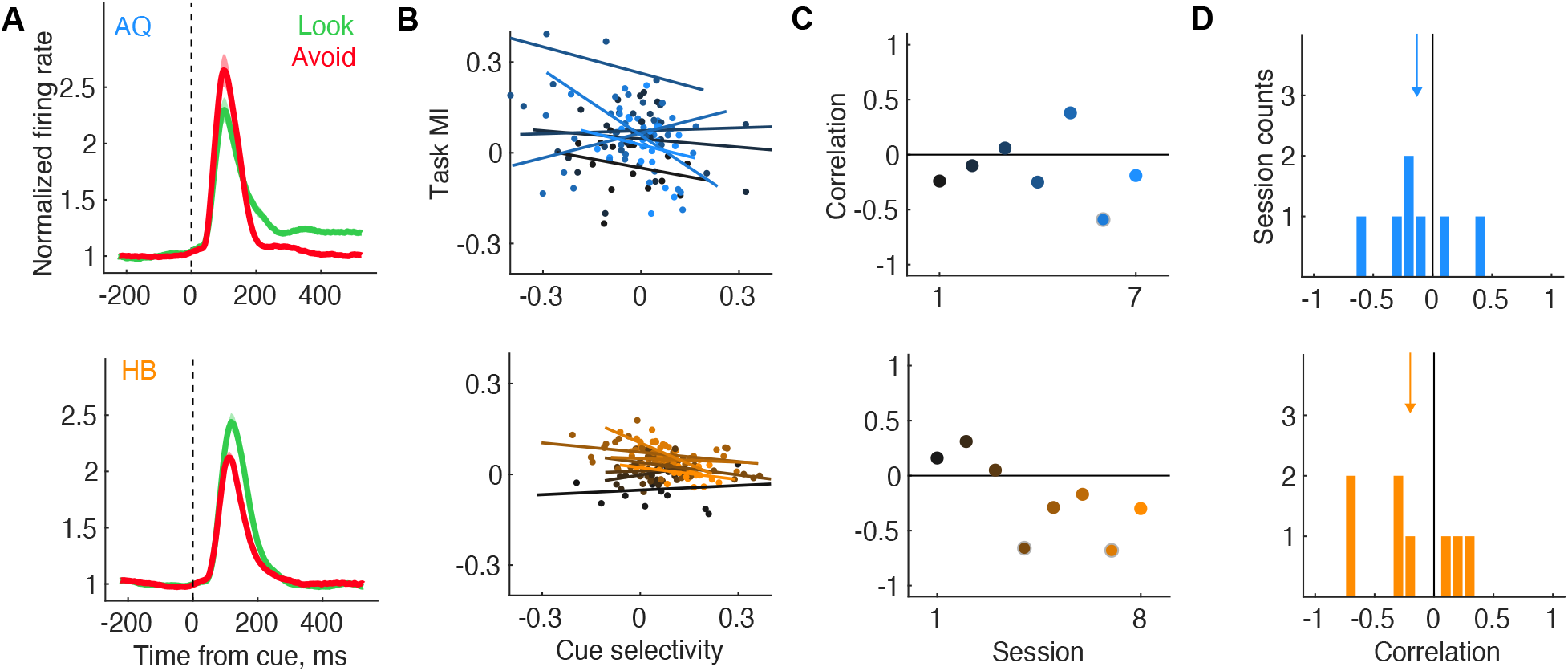
Relationship of pre-cue activity to cue selectivity. **A.** Normalized mean visual responses to the Look and Avoid cues across recordings sessions in the 2 monkeys. The cue stimulus was reversed across monkeys. For AQ, the Look cue was a green square and the avoid cue was a black square. Visual responses were general larger for the (brighter) green square in the 2 monkeys. **B.** Relationship between the cue stimulus selectivity for neurons and their selectivity to the 2 tasks. Stimulus and task selectivity were both measured as modulation indices (MIs)(Methods). Positive task MIs denote greater activity during Look trials, and positive cue selectivity MIs denote greater activity during the presentation of the Look cue. **C.** Correlations between task and cue selectivity MIs in the 2 monkeys. Each set of points and its linear fit shows the correlations for each session. **D.** Distribution of correlations across sessions in the 2 monkeys. If pre-cue activity represents the features of the upcoming cue stimulus, then the two measures should be positively correlated, e.g. neurons selective for the Look cue should exhibit greater activity in the pre-cue period during the Look task. However, we found that these measures were unrelated across sessions in both monkeys. Combined distributions were not significantly different from 0 in either monkey (AQ: *p*=0.21; HB: *p*=0.1) indicating that the precue activity did not predict the responses to the cue stimulus.

**Fig. S6.**
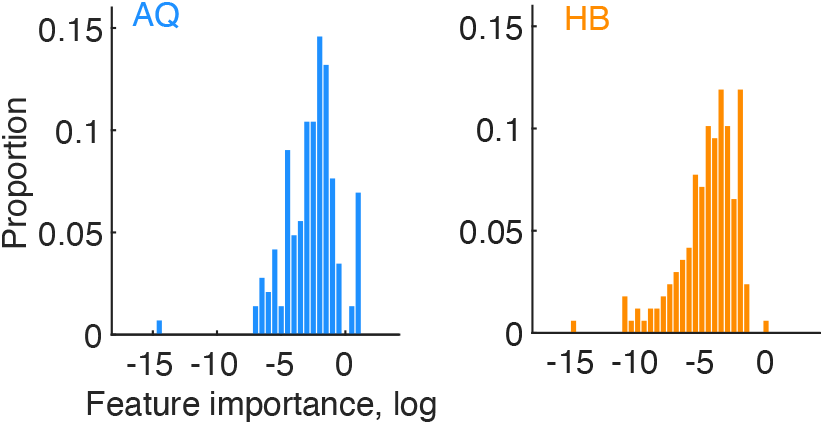
Distribution of feature importance (log units) across sessions in the 2 monkeys.

**Fig. S7.**
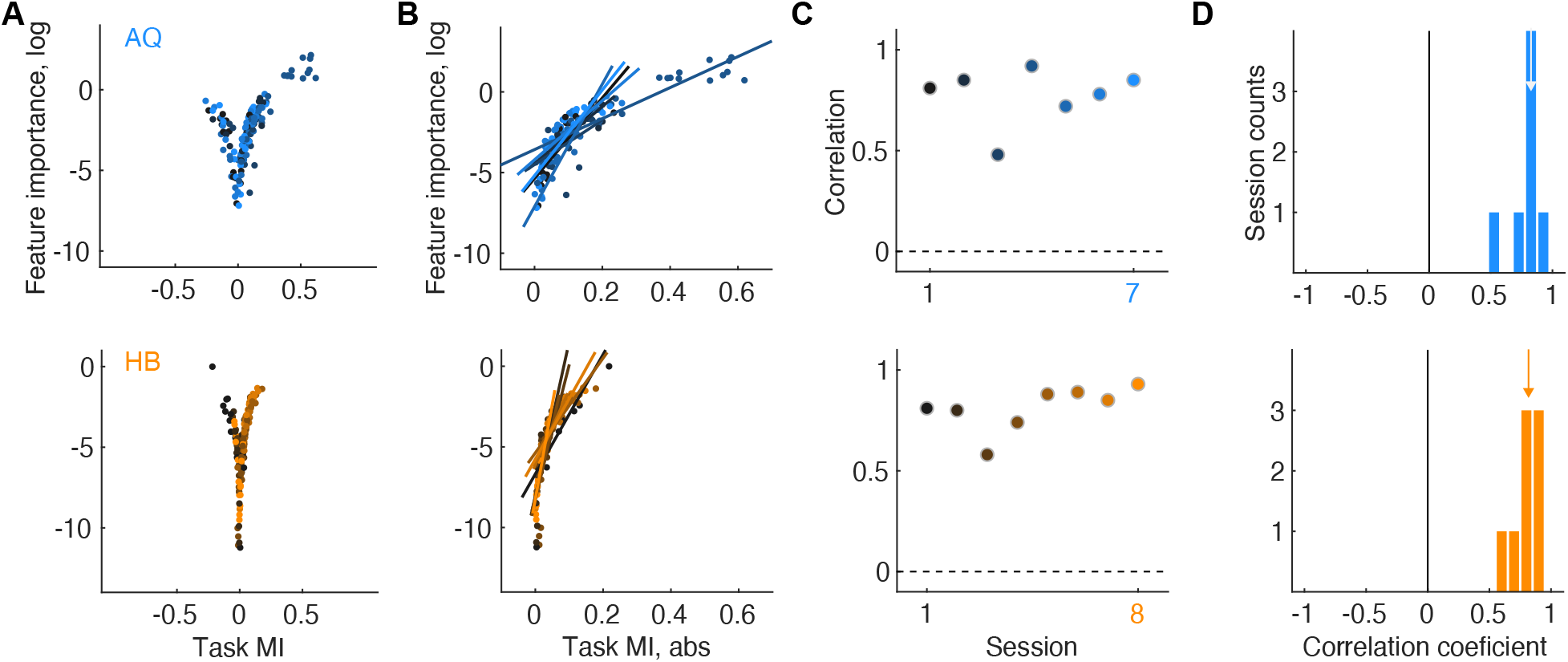
Relationship of feature importance to task modulation. **A.** Plots showing the relationship between neuronal feature importance in the decoder and the task modulation index for the two monkeys (top and bottom). Higher feature importance was associated both with higher and lower modulation indices. **B.** Correlations between feature importance and the absolute task modulation index for each monkey. Each set of points and its linear fit shows the correlations for each session. **C.** Correlation coefficients for each session and monkey. **D.** Distribution of correlation coefficients for both monkeys.

**Fig. S8.**
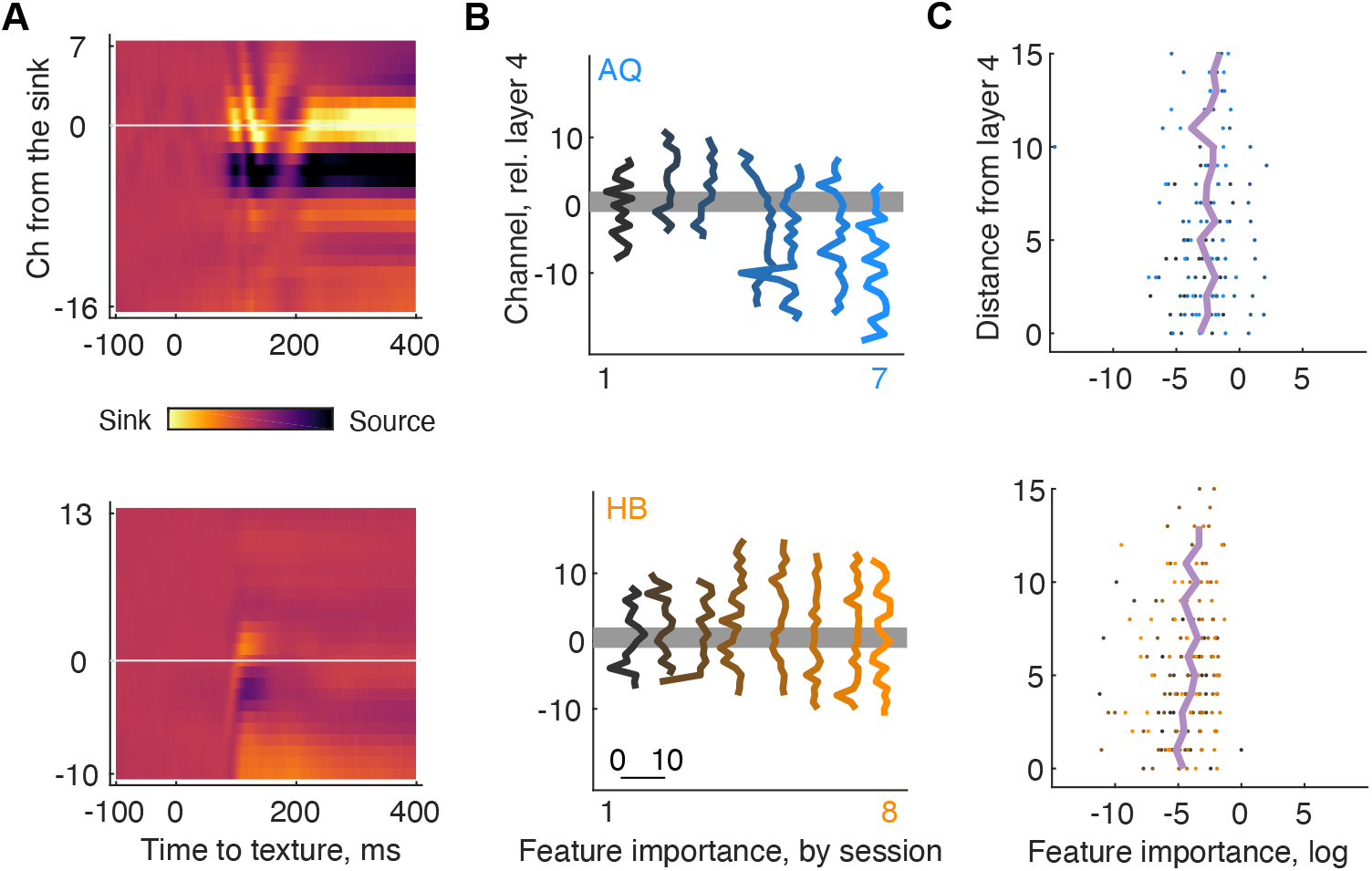
Feature importance as a function of cortical depth. **A.** Example CSD time-course heatmaps for monkey AQ and HB. At time 0 full screen texture appeared (600 ms before cue onset). Input layer (denoted as 0) was determined as the channel with earliest sink response onset. Deeper layers (negative values) and upper layers (positive values). **B.** Decoder weights as a function of depth from the input layer. Data is shown for each individual recording session. **C.** Average decoder weights as a function of absolute distance from input layer. Individual dots represent individual sessions.

## Notes

### Competing Interest Statement

The authors have declared no competing interest.

